# Estimating and interpreting nonlinear receptive fields of sensory responses with deep neural network models

**DOI:** 10.1101/832212

**Authors:** Menoua Keshishian, Hassan Akbari, Bahar Khalighinejad, Jose Herrero, Ashesh D. Mehta, Nima Mesgarani

## Abstract

Sensory processing by neural circuits includes numerous nonlinear transformations that are critical to perception. Our understanding of these nonlinear mechanisms, however, is hindered by the lack of a comprehensive and interpretable computational framework that can model and explain nonlinear signal transformations. Here, we propose a data-driven framework based on deep neural network regression models that can directly learn any nonlinear stimulus-response mapping. A key component of this approach is an analysis method that reformulates the exact function of the trained neural network as a collection of stimulus-dependent linear functions. This locally linear receptive field interpretation of the network function enables straightforward comparison with conventional receptive field models and uncovers nonlinear encoding properties. We demonstrate the efficacy of this framework by predicting the neural responses recorded invasively from the auditory cortex of neurosurgical patients as they listened to speech. Our method significantly improves the prediction accuracy of auditory cortical responses particularly in nonprimary areas. Moreover, interpreting the functions learned by neural networks uncovered three distinct types of nonlinear transformations of speech that varied considerably in primary and nonprimary auditory regions. By combining two desired properties of a computational sensory-response model; the ability to capture arbitrary stimulus-response mappings and maintaining model interpretability, this data-driven method can lead to better neurophysiological models of the sensory processing.

## Introduction

Creating computational models to predict neural responses from the sensory stimuli has been one of the central goals of sensory neuroscience research (1–8). Computational models can be used to form testable hypotheses by predicting the neural response to arbitrary manipulations of stimulus and can provide a way of explaining complex stimulus-response relationships. As such, computational models that provide an intuitive account of how sensory stimuli are encoded in the brain have been critical in discovering the representational and computational principles of sensory cortices (9). One simple yet powerful example of such models is the linear receptive field model, which is commonly used in visual (2,10) and auditory (11,12) neuroscience research. In the auditory domain, linear spectrotemporal receptive field (STRF) (10,12,13) has led to the discovery of neural tuning to various acoustic dimensions, including frequency, response latency, and temporal and spectral modulation (14–16). However, despite the success and ease of interpretability of linear receptive field models, they lack the necessary computational capacity to account for the intrinsic nonlinearities of the sensory processing pathways (17). This shortcoming is particularly problematic in higher cortical areas where stimulus representation becomes increasingly more nonlinear (18,19). Several extensions have been proposed to address the limitations of linear models (20–32) (see (33) for review). These extensions improve the prediction accuracy of neural responses, but this improvement comes at the cost of reduced interpretability of the underlying computation, hence limiting novel insights that can be gained regarding sensory cortical representation. In addition, these methods assumed a specific model structure whose parameters are then fitted to the neural data. This assumed model architecture thus limits the range of the nonlinear transformations that they can account for. This lack of a comprehensive yet interpretable computational framework has hindered our ability to understand the nonlinear signal transformations that are found ubiquitously in the sensory processing pathways (26,34,35).

A general nonlinear modeling framework that has seen great progress in recent years is the multilayer (deep) neural network model (DNN) (36,37). Theses biologically inspired models are universal function approximators (38) and can model any arbitrarily complex input-output relation. Moreover, these data-driven models can learn any form of nonlinearity directly from the data without any explicit assumption or prior knowledge of the nonlinearities. This property makes these models particularly suitable for studying the encoding properties of sensory stimuli in the nervous system (39–41) whose anatomical and functional organization remains largely speculative. A major drawback of DNN models, however, is the difficulty in interpreting the computation that they implement because these models are analytically intractable (42). Thus, despite their success in increasing the accuracy of prediction in stimulus-response mapping, their utility in leading to novel insights into the computation of sensory nervous systems is lacking.

To overcome these challenges, we propose a nonlinear regression framework in which a DNN is used to model sensory receptive fields. An important component of our approach is a novel analysis method that allows for the calculation of the mathematically equivalent function of the trained neural network as a collection of stimulus-dependent, linearized receptive fields. As a result, the exact computation of the neural network model can be explained in a manner similar to that of the commonly used linear receptive field model, which enables direct comparison of the two models. Here, we demonstrate the efficacy of this nonlinear receptive field framework by applying it to neural responses recorded invasively in the human auditory cortex of neurosurgical patients as they listened to natural speech. We demonstrate that not only these models more accurately predict the auditory neural responses, but also uncover distinct nonlinear encoding properties of speech in primary and nonprimary auditory cortical areas. These findings show the critical need for more complete and interpretable encoding models of neural processing, which can lead to better understanding of cortical sensory processing.

## Results

### Neural recordings

To study the nonlinear receptive fields of auditory cortical responses, we used invasive electrocorticography (ECoG) to directly measure neural activity from five neurosurgical patients undergoing treatment for epilepsy surgery. One patient had high-density subdural grid electrodes implanted on the left hemisphere, with coverage primarily over the superior temporal gyrus (STG). All five patients had depth electrodes (stereotactic EEG) with coverage of Heschl’s gyrus (HG) and STG (Fig 1A). While HG and the STG are functionally heterogenous and each contain multiple auditory fields (43–47), HG includes the primary auditory cortex, while the STG is considered mostly a nonprimary auditory cortical area (48). The patients listened to stories spoken by four speakers (two females) with a total duration of 30 minutes. All patients had self-reported normal hearing. To ensure that patients were engaged in the task, we intermittently paused the stories and asked the patients to repeat the last sentence before the pause. Eight separate sentences (40 seconds total) were used as the test data for evaluation of the encoding models, and each sentence was repeated six times in a random order.

**Figure 1.**
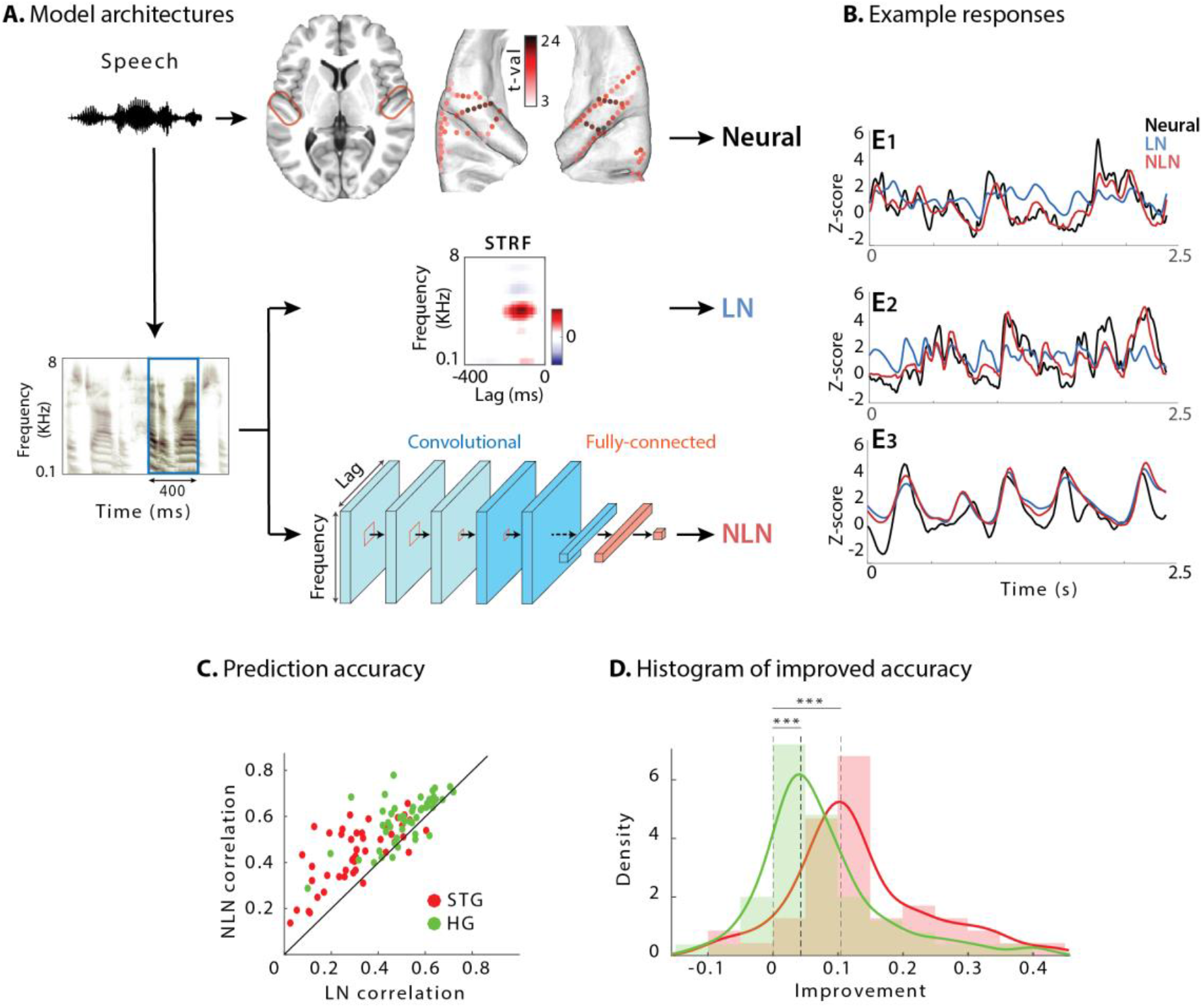
Predicting neural responses using linear and nonlinear regression models. (A) Neural responses to speech were recorded invasively from neurosurgical patients as they listened to speech. The brain plot shows electrode locations and t-value of the difference between the average response of a neural site to speech versus silence. The neural responses are predicted from the stimulus using a linear spectrotemporal receptive field model (LN) and a nonlinear convolutional neural network model (NLN). Input to both models is a sliding time-frequency window with 400 ms duration. (B) Actual and predicted responses of three example sites using the LN and NLN models. (C) Prediction accuracy of neural responses from the LN and NLN models for sites in STG and HG. (D) Distribution of improved accuracy of predicted responses over the linear model for all sites in HG and STG.

We used the envelope of the high-gamma (70-150 Hz) band of the recorded signal as our neural response measure, which correlates with the neural firing in the proximity of electrodes (49,50). The high-gamma envelope was extracted by first filtering neural signals with a bandpass filter and then calculating the Hilbert envelope. We restricted our analysis to speech-responsive electrodes, which were selected using a t-test between the average response of each electrode to speech stimuli versus the response in silence (t-value > 2). This criterion resulted in 50 electrodes in HG and 47 in STG. Electrode locations are plotted in Fig 1A on the average FreeSurfer brain (51) where the colors indicate speech responsiveness (t-values).

We used the auditory spectrogram of speech utterances as the acoustic representation of the stimulus. Auditory spectrograms were calculated using a model of the peripheral auditory system that estimates a time-frequency representation of the acoustic signal on a tonotopic axis (52). The speech materials were split into three subsets for fitting the models – training, validation, and test. The test was obtained by averaging the neural responses to the repeated sentences to reduce the effect of neural noise on model comparisons. The remainder of the data was split between training and validation subsets (97% and 3% respectively). There was no stimulus overlap between the three subsets.

### Linear and nonlinear encoding models

For each neural site, we fit one linear (LN) and one nonlinear (NLN) regression model to predict its activity from the auditory spectrogram of the stimulus (Fig 1A). The input to the regression models was a sliding window with a duration of 400 ms with 10 ms steps, which was found by maximizing the prediction accuracy (Supplementary Fig 1). The linear encoding model was a conventional STRF that calculates the linear mapping from stimulus spectrotemporal features to the neural response (10). The nonlinear regression model was implemented using a deep convolutional neural network (CNN (53)) consisting of two stages: a feature extraction network and a feature summation network. This framework, commonly used for nonlinear regression (54–56), consists of extracting a high-dimensional representation of the input (feature extraction), followed by a feature summation network to predict neural responses. The feature extraction network comprises three convolutional layers each with eight 3×3 2D convolutional kernels, followed by two convolutional kernels with 1×1 kernels to reduce the dimensionality of the representation, thus decreasing the number of model parameters. The feature summation stage was a two-layer fully-connected network with 32 nodes in the hidden layer and a single output node. All hyperparameters of the network were determined by optimizing the prediction accuracy (Supplementary Fig 2). All hidden layers had rectified linear unit (ReLU) nonlinearity (57), and the output node was linear. A combination of mean-squared error and Pearson correlation was used as the training loss function (see Methods).

### Predicting neural responses using linear and nonlinear encoding models

Examples of actual and predicted neural responses from the LN and NLN models are shown in Fig 1B for three sample electrodes. We examined the nonlinearity of each neural site by comparing the prediction accuracy of LN and NLN models. As Fig 1B shows, the NLN predictions (red) are more similar to the actual responses (black) compared to LN predictions (blue). This observation confirms that the NLN model can capture the variations in the neural responses to speech stimuli more accurately. To quantify this improvement across all sites, we calculated the noise-adjusted R-squared value (58) (see Methods) between the predicted and actual neural responses for each model. The scatter plot in Fig 1C shows the comparison of these values obtained for LN and NLN models for each electrode, where the electrodes are colored by their respective brain region. Fig 1C shows higher accuracy for NLN predictions compared to LN predictions for the majority of electrodes (87 out of 97). Even though the overall prediction accuracy is higher for HG electrodes than ones in STG, the absolute improved prediction of NLN over LN is significantly higher for STG electrodes (Fig 1D; p < 0.003, one-tailed t-test). This higher improvement in STG electrodes reveals a larger degree of nonlinearity in the encoding of speech in the STG compared to HG.

### Interpreting the nonlinear receptive field learned by DNNs

The previous analysis demonstrates the superior ability of the NLN model to predict the cortical representation of speech particularly in higher order areas, but it does not show what types of nonlinear computation results in improved prediction accuracy. To explain the mapping learned by the NLN model, we developed an analysis framework that finds the mathematical equivalent linear function that the neural network applies to each instance of the stimulus. This equivalent function is found by estimating the derivative of the network output with respect to its input (i.e., the data Jacobian matrix (59)). We refer to this equivalent function as the locally linear receptive field (LLRF). The LLRF can be considered a STRF whose spectrotemporal tuning depends on every instance of the stimulus. As a result, the linear weighting function of the NLN model that is applied to each stimulus instance can be visualized in a manner similar to that of the STRF (see Supplementary Videos 1 and 2).

Finding the LLRF is particularly straightforward for a neural network with rectified linear unit nodes (ReLU), because ReLU networks implement piecewise linear functions (Fig 2C). A rectified linear node is inactive when the sum of its inputs is negative or is active and behaves like a linear function when the sum of its inputs is positive (Fig 2A). Obtaining a linear equivalent of such a DNN for a given stimulus consists of first removing the weights that connect to the inactive nodes in the network (Fig 2B) and replacing the active nodes with identity functions. Next, the remaining weights of each layer are multiplied to calculate the overall linear weighting function applied to the stimulus. Because the resulting weighting vector has the same dimension as the input to the network, it can be visualized as a multiplicative template similar to STRF (Fig 2B). The mathematical derivation of LLRF is shown in the Methods.

**Figure 2.**
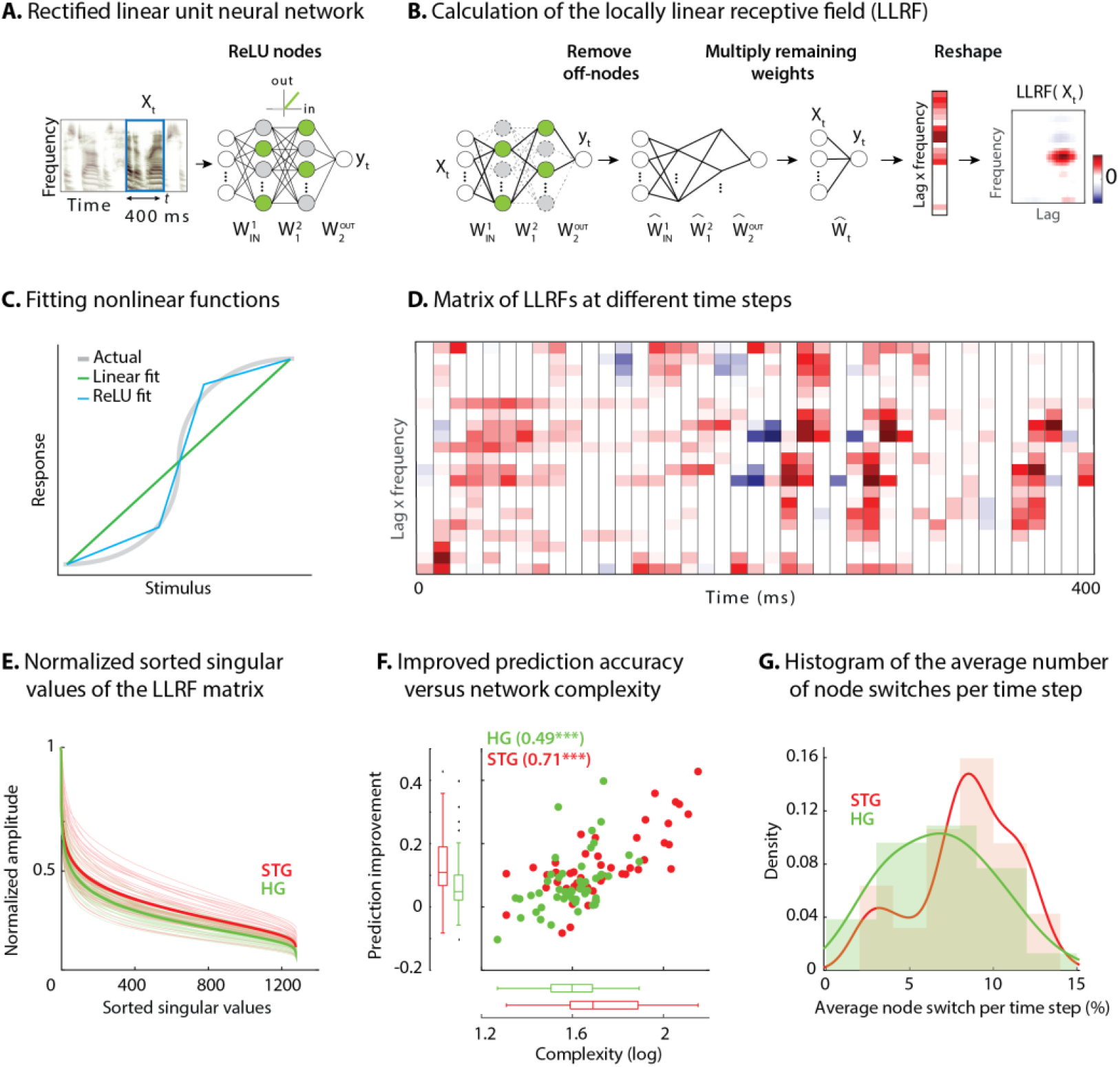
Calculating the stimulus-dependent linearized spectrotemporal receptive field (LLRF) (A) Activation of nodes in a neural network with rectified linear (ReLU) nonlinearity for the stimulus at time *t*. (B) Calculating the stimulus-dependent locally linear receptive field (LLRF) for input instance *X_t_* by first removing all inactive nodes from the network and replacing the active nodes with the identity function. The LLRF is then computed by multiplying the remaining network weights. Reshaping the resulting weight matrix expresses the LLRF in the same dimensions as the input stimulus and can be interpreted as a multiplicative template applied to the input. (C) Comparison of piecewise linear (rectified linear neural network) and linear (STRF) approximations of a nonlinear function. (D) LLRF vectors (columns) shown for 40 samples of the stimulus. Only a limited number of lags and frequencies are shown at each time step to assist visualization. (E) Normalized sorted singular values of the LLRF matrix show higher diversity of the learned linear function in STG sites than in HG sites. The bold lines are the averages of sites in the STG and in HG. The complexity of a network is defined as the sum of the sorted singular values. (F) Comparison of network complexity and the improved prediction accuracy over the linear model in STG and HG areas. (G) Histogram of the average number of switches from on/off to off/on states at each time step for the neural sites in the STG and HG.

An example of LLRF at different time points for one electrode is shown in Fig 2D where each column is the vectorized LLRF (time lag by frequency) that is applied to the stimulus at that time point (for better visibility only part of the actual matrix is shown). This matrix contains all the variations of the receptive fields that the network applies to the stimulus at different time points. On one extreme, a network could apply a fixed receptive field to the stimulus at all times (similar to the LN model) for which the columns of the matrix in Fig 2D will all be identical. At the other extreme, a network can learn a unique function for each instance of the stimulus for which the columns of the matrix in Fig 2D will all be different functions. Because a more nonlinear function results in a higher number of piecewise linear regions (60) (Fig 2C), the diversity of the functions in the lag-frequency by time matrix (columns in Fig 2D) indicates the degree of nonlinearity of the function that the network has learned. To quantify this network nonlinearity, we used singular-value decomposition (61) of the lag-frequency by time matrix. Each singular value indicates the variance in its corresponding dimension; therefore, the steepness of the sorted singular values is inversely proportional to the diversity of the learned functions. The sorted normalized singular values for all electrodes in HG and STG are shown in Fig 2E, demonstrating that the neural network models learn considerably more diverse functions for STG electrodes. This result confirms the increase in nonlinearity observed earlier in STG electrodes compared to HG electrodes.

### Complexity of the nonlinear receptive field

We defined the complexity of the NLN model as the sum of the normalized singular values of the lag-frequency by time matrix (Fig 2D). The complexity for all electrodes in HG and STG is shown in Fig 2F and is compared against the improved prediction accuracy of the NLN over the LN model. Fig 2F shows a significantly higher complexity for STG electrodes than for HG electrodes (p < 0.001, t-test), which also correlates with the prediction improvement of each electrode over the linear model (r = 0.66, p < 0.001). Alternatively, a separate metric to measure the degree of network nonlinearity is the average number of nodes that switch between active and inactive states at each time step. The histogram of the average switches for HG and STG electrodes shows significantly higher values for STG electrodes (Fig 2G; p < 0.01, one-tailed t-test). This observation validates the finding that the larger improvement of prediction accuracy in STG is due to the implementation of a more diverse set of linear templates, whereas the HG electrodes require a smaller number of linear functions to accurately capture the stimulus response relationships in this area. Importantly, these results are not dependent on network parameters and network initialization. While the hyperparameters and the training of the network can change the internal implementation of the input-output function, the function itself is robust and remains unchanged (Methods and Supplementary Fig 3). Furthermore, the linearized functions averaged over all samples closely resemble the STRF for the corresponding electrode, with the similarity decreasing with the complexity of the neural site. (Supplementary Fig 4).

### Identifying various types of nonlinear receptive field properties

We showed that the nonlinear function of the NLN model can be expressed as a collection of linearized functions. To investigate the properties of these linearized functions learned by the models for various neural sites, we visually inspected the LLRFs and observed three general types of variations over time, which we refer to as: I) gain change, II) temporal hold, and III) shape change. We describe and quantify each of these three nonlinear computations in this section using three example electrodes that exhibit each of these types more prominently (Supplementary Video 3). The STRFs for the three examples sites are shown in Fig 3A.

**Figure 3.**
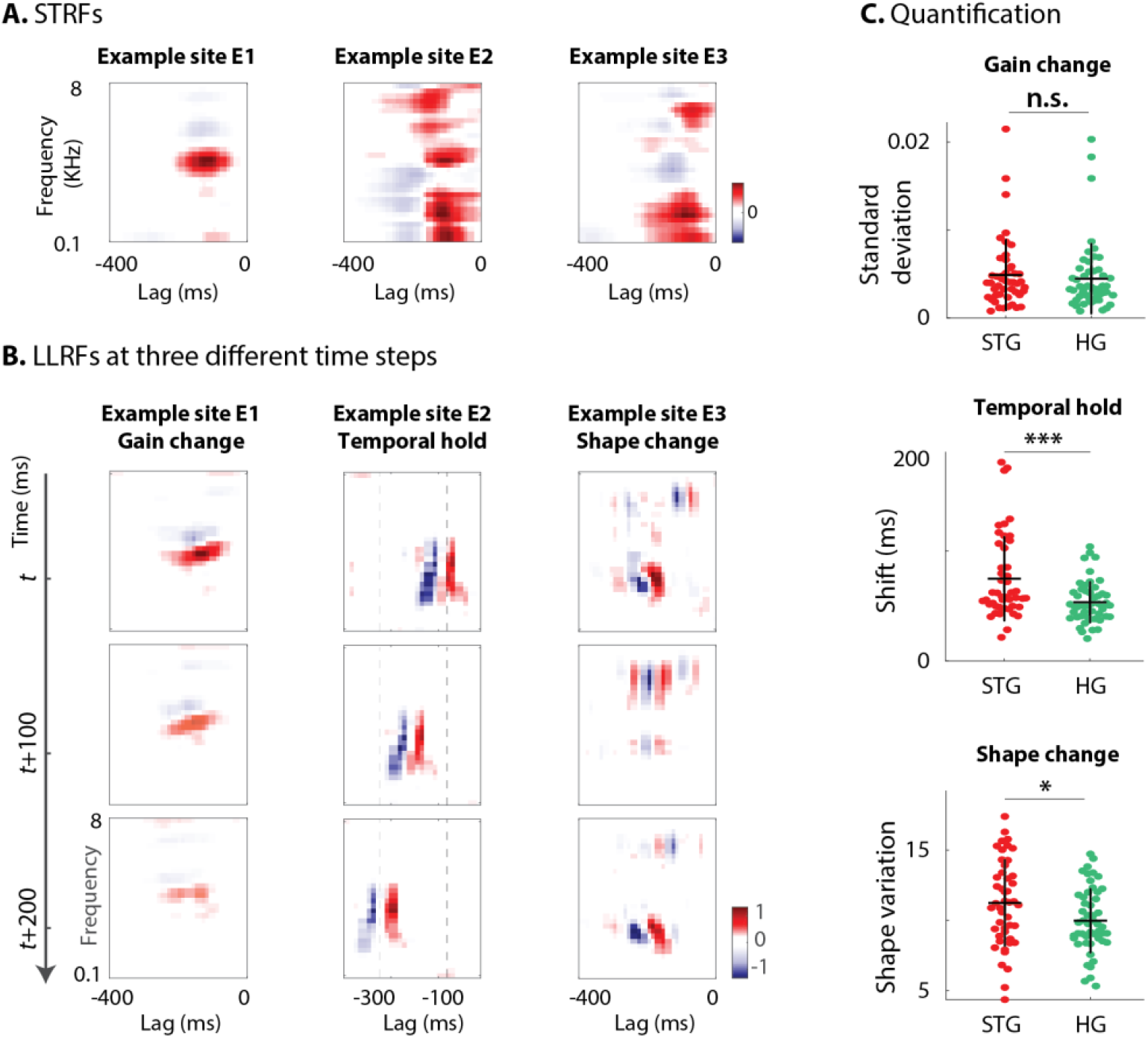
Characterizing the types of LLRF variations. Three types of LLRF variations over time for three example sites that exhibit each of these types more prominently. (A) The STRF of the three example sites. (B) Example site E1: Gain change of LLRF, shown as the time-varying magnitude of the LLRF at three different time points. Although the overall shape of the spectrotemporal receptive field is the same, its magnitude varies across time. E2: Temporal hold property of an LLRF, seen as the tuning of this site to a fixed spectrotemporal feature but with shifted latency (lag) in consecutive time frames. E3: Shape change property of LLRF, seen as the change in the spectrotemporal tuning pattern for different instances of the stimulus. (C) From top to bottom, quantification of gain change, temporal hold, and shape change for each site in STG and HG. Temporal hold and shape change show significantly higher values in the STG.

The first and simplest type of LLRF change is gain change, which refers to the time-varying magnitude of the LLRF. This effect is shown for one example site (E1) at three different time points in Fig 3B. Although the overall shape of the spectrotemporal filter applied to the stimulus at these three time points is not different, the magnitude of the function varies considerably over time. This variation in the gain of the stimulus-response function that is learned by the DNN may reflect the nonlinear adaptation of neural responses to the acoustic stimulus (34,62,63). We quantified the degree of gain change for each site by calculating the standard deviation of the LLRF magnitude over the stimulus duration.

The second type of LLRF change is temporal hold, which refers to the tuning of a site to a fixed spectrotemporal feature but with shifted latency in consecutive time frames (Fig 3B). This particular tuning nonlinearity results in a sustained response to arbitrarily fast spectrotemporal features, hence resembling a short-term memory of that feature. An example of temporal hold for a site (E2) is shown in Fig 3B where the three plots show the LLRF of this site at three consecutive time steps. Even though the overall shape of the spectrotemporal feature tuning remains the same, the latency (lag) of the feature shifts over time with the stimulus. This nonlinear property decouples the duration of the response to an acoustic feature from the temporal resolution of that feature. For example, temporal hold could allow a network to model a slow response to fast acoustic features. This computation cannot be done with a linear operation because a linear increase in the analysis time scale inevitably results in the loss of temporal resolution, as shown in the STRF of site E2 in Fig 3A. We quantified the temporal hold for each site using the cross correlation of LLRFs at each time point with the LLRFs in the following time steps. Then, we calculated the number of frames for which the shifted spectrotemporal feature appears in the LLRF. The average duration of the shifted spectrotemporal features is defined as the temporal hold for each site (see Supplementary Fig 5 for graphic illustration).

The last dimension of LLRF variation is shape change, which refers to a change in the spectrotemporal tuning of a site across stimuli. Intuitively, a model that implements a more nonlinear function will have a larger number of piecewise linear regions, each exhibiting a different LLRF shape. An example of shape change for a site (E3) is shown in Fig 3B, showing three different spectrotemporal patterns at these three different time points. To quantify the degree of change in the shape of the LLRF for a site, we repeated the calculation of network complexity but after normalizing and removing the effect of gain change and temporal hold from the LLRFs. Therefore, shape change is defined as the sum of the normalized singular values of the gain- and shift-corrected lag-frequency by time matrix (Fig 2D). Thus, the shape change indicates the remaining complexity of the LLRF function that is not due to gain change or temporal hold. This stimulus-dependent change in the spectrotemporal tuning of sites reflects a nonlinearity that appears as the sum of all possible shapes in the STRF, as shown in Fig 3B.

The distribution of the three nonlinearity types defined here across all neural sites in HG and STG areas are shown in Fig 3C. Looking at these distributions can give us a better understanding of the shared nonlinear properties among neural populations of each region. The degree of gain change for electrodes in HG and STG areas spans a wide range. However, we did not observe a significant difference between the gain change values in STG and HG sites (p > 0.6, t-test), suggesting a similar degree of adaptive response in HG and STG sites. The distribution of temporal hold values for all sites in the STG and HG shows significantly larger values in STG sites (Fig 3C, p < 0.001). This increased temporal hold is consistent with the previous findings showing increased temporal integration in higher auditory cortical areas (64,65), which allows the stimulus information to be combined across longer time scales. Finally, the average of the shape change values is significantly higher in STG sites than in HG (p < 0.05), demonstrating that STG sites have more nonlinear encoding, which requires more diverse receptive field templates than HG sites.

### Finding subtypes of receptive fields

As explained in the shape change nonlinearity, the NLN model may learn several subtypes of receptive fields for a neutral site. To further investigate the subtypes of receptive fields that the NLN model learns for each site, we used the k-means algorithm (66) to cluster shift-corrected LLRFs based on their correlation similarity. The optimal number of clusters for each site was determined using the gap statistic method (67). The optimal number of clusters across all sites differed from 1 to 6; the majority of sites, however, contained only one main cluster (84.9% of sites; mean number of clusters = 1.43 ± 1.26 SD). Fig 4 shows the LLRF clustering analysis for two example sites, where for each cluster the average LLRF and the average auditory stimulus for the time points that have LLRFs belonging to that cluster are shown. The average LLRFs in Fig 4 show two distinct receptive field shapes that the NLN models apply to the stimulus at different time points. In addition, the average spectrograms demonstrate the distinct time-frequency power in the stimuli that caused the model to choose the corresponding template.

**Figure 4.**
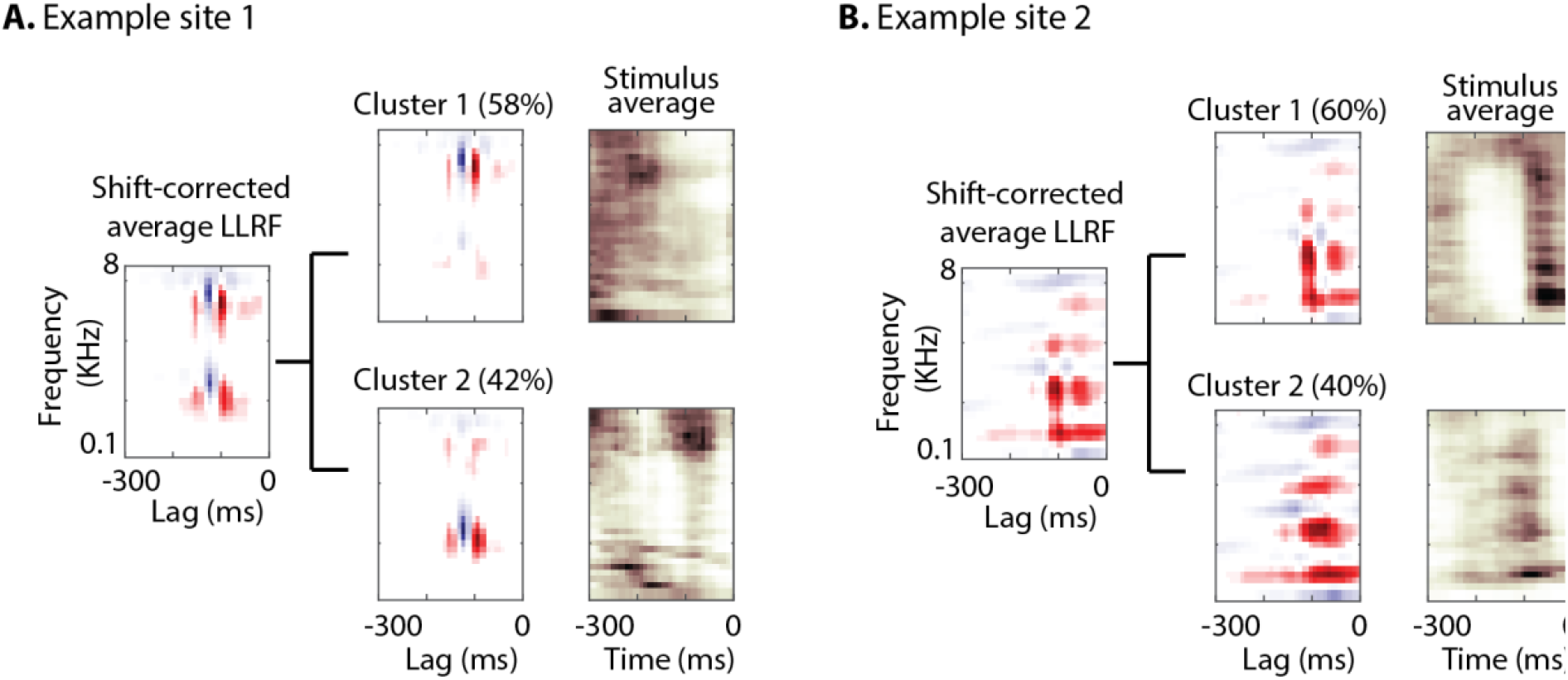
Characterizing the spectrotemporal tuning of electrodes with clustering. (A, B) The average shift-corrected LLRF for two example sites and average k-means clustered LLRFs based on the similarity of LLRFs (correlation distance). Each cluster shows tuning to a distinct spectrotemporal feature. The average LLRF shows the sum of these different features. The average spectrograms over the time points at which each cluster LLRF was selected by the network demonstrate the distinct time-frequency pattern in the stimuli that caused the network to choose the corresponding template.

### Contribution of nonlinear variations to network complexity and prediction improvement

So far, we have defined and quantified three types of nonlinear computation for each neural site – gain change, temporal hold, and shape change – resulting in three numbers describing the stimulus-response nonlinearity of the corresponding neural population. Next, we examined how much of the network complexity (Fig 2F) and improved accuracy over a linear model (Fig 1D) can be accounted for by these three parameters. We used linear regression to calculate the complexity and prediction improvement for each site from the gain change, temporal hold, and shape change parameters. The predicted and actual complexity of the models are shown in Supplementary Fig 6. The high correlation value (r = 0.92, p << 1) confirms the efficacy of these three parameters to characterize the complexity of the stimulus-response mapping across sites. Moreover, the main effects of the regression (68) shown in Fig 5A suggest a significant contribution from all three parameters to the overall complexity of LLRFs in both HG and STG.The high correlation values between the actual and predicted improved accuracy (Supplementary Fig 6; r = 0.72, p < 0.001) show that these three parameters also largely predict stimulus-response nonlinearity. The contribution of each nonlinear factor in predicting the improved prediction, however, is different between the HG and STG areas, as shown by the main effects of the regression in Fig 5B. All three factors contribute to the improved accuracy in STG sites, while only the gain change is a good predictor of the improvement in HG sites. This result reiterates the encoding distinctions we observed in HG and STG sites where temporal hold and shape change were significantly higher in STG than HG (Fig 3C). Notably, the three types of LLRF variations are not independent of each other and covary considerably (Supplementary Fig 7). Together, these results demonstrate how our proposed nonlinear encoding method can lead to a comprehensive, intuitive way of studying nonlinear mechanisms in sensory neural processing by utilizing deep neural networks.

**Figure 5.**
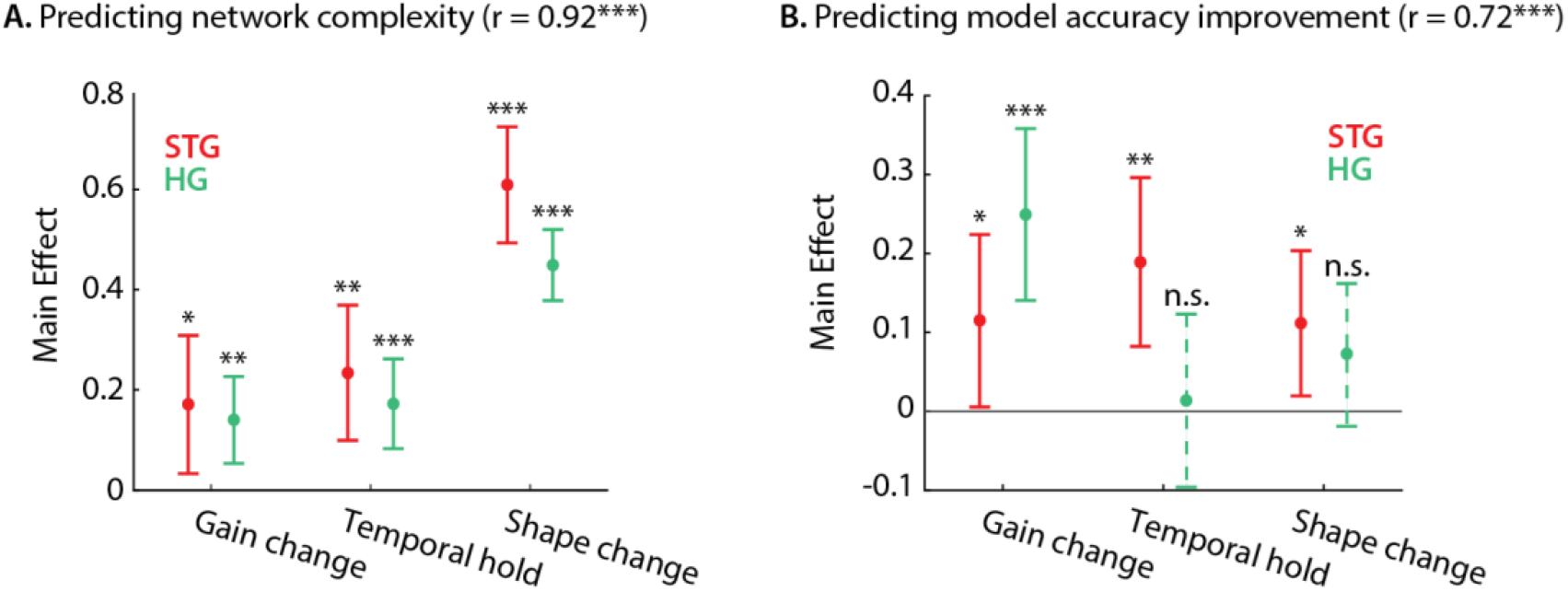
Contribution of LLRF variations to network complexity and prediction improvement. (A) Predicting the complexity of the neural networks from gain change, temporal hold, and shape change using a linear regression model. (B) The main effect of regression analysis shows the significant contribution of all nonlinear parameters in predicting the network complexity in both STG and in HG sites. (C) Predicting the improved accuracy of neural networks over the linear model from the three nonlinear parameters for each site in the STG and HG. (D) The main effect of the regression analysis shows the different contribution of the threes nonlinear parameters in predicting the improved accuracy over the linear model in different auditory cortical areas.

## Discussion

We propose a general nonlinear regression framework to model and interpret any complex stimulus-response mapping in sensory neural responses. This data-driven method provides a unified framework that can discover and model various types of nonlinear transformations without needing any prior assumption about the nature of nonlinearities. In addition, the function learned by the network can be interpreted as a collection of linear receptive fields from which the network chooses for different instances of the stimulus. We demonstrated how this method can be applied to auditory cortical responses in humans to discover novel types of nonlinear transformation of speech signal, therefore extending our knowledge of the nonlinear cortical computations beyond what can be explained with previous models. Together, our results showed three distinct nonlinear properties which largely account for the complexity of the neural network function and could predict the improved prediction accuracy over the linear model. Extracting these nonlinear components and incorporating them in simpler models such as the STRF can systematically test the contribution and interaction of these nonlinear aspects more carefully. However, such simplification and abstraction proved nontrivial in our data because the different types of variation we describe in this paper are not independent of each other and covary considerably.

The increasing nonlinearity of the stimulus-response mapping throughout the sensory pathways (64,69) highlights the critical need for nonlinear computational models of sensory neural processing, particularly in higher cortical areas. These important nonlinear transformations include nonmonotonic and nonlinear stimulus-response functions (70), time-varying response properties such as stimulus adaptation and gain control (34,62,63,71), and nonlinear interaction of stimulus dimensions and time-varying stimulus encoding (72). These nonlinear effects are instrumental in creating robust perception, which requires the formation of invariant perceptual categories from a highly variable stimulus (73–79). The previous research on extending simple receptive field models that have tried to address the linear system limitations include generalized linear models (20), linear-nonlinear (LN) models (21–23), input nonlinearity models (24,80), gain control models (26–28), context-dependent encoding models, and LNLN cascade models (29–32) (see (33) for review). Even though all these models improve the prediction accuracy of neural responses, this improvement comes at the cost of reduced interpretability of the computation. For example, the multifilter extensions of the auditory STRF (21–23,29–32) lead to nonlinear interactions of multiple linear models, which is considerably less interpretable than the STRF model. Our approach extends the previous methods by significantly improving the prediction accuracy over the linear model and at the same time, remaining highly interpretable.

The computational framework we propose to explain the neural network function (81) can be used in any feedforward neural network model with any number of layers and nodes, such as in fully connected networks, locally connected networks (82), or CNNs (83). Nonetheless, one limitation of the LLRF method is that it cannot be used for neural networks with recurrent connections. Because feedforward models use a fixed duration of the signal as the input, the range of the temporal dependencies and contextual effects that can be captured with these models is limited. Nevertheless, sensory signals such as speech have long-range temporal dependencies for which recurrent networks may provide a better fit. Although we did not find a significant difference between the prediction accuracy of feedforward and recurrent neural networks in our data (Supplementary Fig 8), the recent extensions of the feedforward architecture, such as dilated convolution (84) or temporal convolutional networks (85), can implement receptive fields that extend over long durations. Our proposed LLRF method would seamlessly generalize to these architectures, which can serve as an alternative to recurrent neural networks when modeling the long-term dependencies of the stimulus is crucial.

In summary, our proposed framework combines two desired properties of a computational sensory-response model; the ability to capture arbitrary stimulus-response mappings and maintaining model interpretability. We showed that this data-driven method reveals novel nonlinear properties of cortical representation of speech in the human brain which provides an example for how it can be used to create more complete neurophysiological models of sensory processing in the brain.

## Online Methods

### Participants and neural recording

Five patients with pharmacoresistant focal epilepsy were included in this study. All patients underwent chronic intracranial encephalography (iEEG) monitoring at Northshore University Hospital to identify epileptogenic foci in the brain for later removal. Four patients were implanted with stereo-electroencephalographic (sEEG) depth arrays only and one with both depth electrodes and a high-density grid (PMT, Chanhassen, MN, USA). Electrodes showing any sign of abnormal epileptiform discharges, as identified in the epileptologists’ clinical reports, were excluded from the analysis. All included iEEG time series were manually inspected for signal quality and were free from interictal spikes. All research protocols were approved and monitored by the institutional review board at the Feinstein Institute for Medical Research, and informed written consent to participate in research studies was obtained from each patient before electrode implantation.

iEEG signals were acquired continuously at 3 kHz per channel (16-bit precision, range ± 8 mV, DC) with a data acquisition module (Tucker-Davis Technologies, Alachua, FL, USA). Either subdural or skull electrodes were used as references, as dictated by recording quality at the bedside after online visualization of the spectrogram of the signal. Speech signals were recorded simultaneously with the iEEG for subsequent offline analysis. The envelope of the high-gamma response (75-150 Hz) was extracted by first filtering neural signals with a bandpass filter and then using the Hilbert transform to calculate the envelope. The high-gamma responses were z-scored and resampled to 100 Hz.

### Brain maps

Electrode positions were mapped to brain anatomy using registration of the postimplant computed tomography (CT) to the preimplant MRI via the postop MRI. After coregistration, electrodes were identified on the postimplantation CT scan using BioImage Suite. Following coregistration, subdural grid and strip electrodes were snapped to the closest point on the reconstructed brain surface of the preimplantation MRI. We used FreeSurfer automated cortical parcellation (51) to identify the anatomical regions in which each electrode contact was located with a resolution of approximately 3 mm (the maximum parcellation error of a given electrode to a parcellated area was < 5 voxels/mm). We used Destrieux parcellation, which provides higher specificity in the ventral and lateral aspects of the medial lobe. Automated parcellation results for each electrode were closely inspected by a neurosurgeon using the patient’s coregistered postimplant MRI.

### Stimulus

Speech materials consisted of continuous speech stories spoken by four speakers (2 male and 2 female). The duration of the stimulus was 30 minutes and was sampled at 11025 Hz. Eight sentences (40 seconds) were used for testing and presented to the patients six times to improve the signal-to-noise ratio. The 30-minute data were split into two segments for training (30 minutes) and validation (50 seconds). There was no overlap between the training, test, and validation sets. The input to the regression models was a sliding window of 400 ms (40 timesteps), which was chosen to optimize prediction accuracy (Supplementary Fig 1). The windowing stride was set to one to maintain the same final sampling rate, and as a result, the two consecutive input vectors to the regression models overlapped at 39 time points.

### Acoustic representation

An auditory spectrogram representation of speech was calculated from a model of the peripheral auditory system (52). This model consists of the following stages: 1) a cochlear filter bank consisting of 128 constant-Q filters equally spaced on a logarithmic axis, 2) a hair cell stage consisting of a low pass filter and a nonlinear compression function, and 3) a lateral inhibitory network consisting of a first-order derivative along the spectral axis. Finally, the envelope of each frequency band was calculated to obtain a time-frequency representation simulating the pattern of activity on the auditory nerve. The final spectrogram has a sampling frequency of 100 Hz. The spectral dimension was downsampled from 128 frequency channels to 32 channels to reduce the model complexity.

### Calculating spectrotemporal receptive fields (STRFs)

Linear STRF models were fitted using the STRFlab MATLAB toolbox (86,87). For each electrode, a causal model was trained to predict the neural response at each time point from the past 400 ms of stimulus. The optimal model sparsity and regularization parameters were chosen by maximizing the mutual information between the actual and predicted responses for each electrode.

### DNN architecture

We designed a two-stage DNN consisting of feature extraction and feature summation modules (Fig 1A). In this framework, a high-dimensional representation of the input is first calculated (feature extraction network), and this representation is then used to regress the output of the model (feature summation network). The feature extraction stage consists of three convolutional layers with eight 3×3 2D convolutional kernels each, followed by a convolutional layer with four 1×1 kernels to reduce the dimensionality of the representation. The output of this stage is the input to another convolutional layer with a single 1×1 kernel. A 1×1 kernel that is applied to an input with *N* channels has 1×1×*N* parameters, and its output is a linear combination of the input channels, thus reducing the dimension of the latent variables and, consequently, the number of trainable parameters. The feature summation stage is a two-layer fully connected network with a hidden layer of 32 nodes, followed by an output layer with a single node. All layers except the output layer have L2 regularization, dropout (88), and ReLU (57) activations. The output layer has regularization and linear activation. The parameters of the model, including the number of layers, the size of the convolutional kernels, and the number of fully connected nodes, were found by optimizing the prediction accuracy (Supplementary Fig 2).

### DNN training and cross-validation

The networks were implemented in Keras using the TensorFlow backend. A separate network was trained for each electrode. Kernel weight initializations were performed using a method specifically developed for DNNs with rectified linear nonlinearities (89) for faster convergence. We used ReLU nonlinearities for all layers except the last layer, and dropouts with *p* = 0.3 for the convolutional layers and *p* = 0.4 for the first fully connected layer were used to maximize prediction accuracy. The convolutional layers had strides of one, and their inputs were padded with zeros such that the layer’s output would have the same dimension as the input. We applied an *L2* penalty (with a multiplier weight set to 0.001) to the weights of all the layers. Each training epoch had a batch size of 128, and optimization was performed using Adam with an initial learning rate of 0.0001. Networks were trained with a maximum of 30 epochs, with early stopping when the validation loss did not decrease for five consecutive epochs. The weights that resulted in the best validation loss during all training epochs were chosen as the final weights. The loss function was a linear combination of the MSE and Pearson’s correlation coefficient:

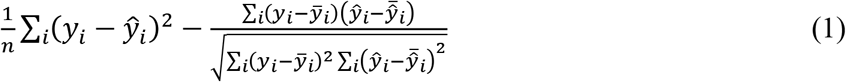

in which *y* is the high-gamma envelope of the recorded neural data from a given electrode, and *ŷ* is the predicted response of the neural network.

### Evaluating model performance (test-retest)

To account for the variations in neural responses that are not due to the acoustic stimulus, we repeated the test stimulus six times to more accurately measure the explainable variance. To obtain a better measure of the model’s goodness of fit, we used a noise-corrected adjusted R-squared value instead of the simple correlation. Having *n* responses to the same stimulus, *r*_1_ to *r_n_*, we defined *r_o_* and *r_e_* as the averages of odd and even numbered trials. Then, we calculated the noise-corrected correlation according to equation (2), where *p* is the predicted response of a model, *C*_1_ is the average correlation of the predicted and actual responses, *C*_2_ is the correlation between response groups, and *C* is our reported R-squared of the noise-corrected Pearson correlation.

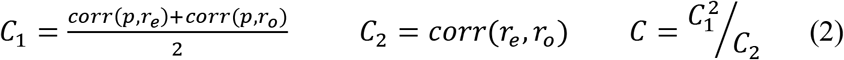

### Computing locally linear receptive fields for convolutional neural networks

The first step for calculating the locally linear receptive field (LLRF) of a CNN consists of converting the CNN into a multilayer perceptron (MLP) (Supplementary Fig 9) because calculating LLRFs for an MLP is more straightforward. To achieve this task, we must first convert each convolutional layer to its equivalent locally connected layer, which is essentially a sparse fully connected layer. To do so, we find the equivalent matrix *W* for convolutional kernels *K*_1_ – *K_l_*, where *K_i_* is the *i*-th kernel of the convolutional layer. Transforming all layers of the CNN into fully connected layers results in an MLP network. The input and output tensors of the fully connected layers have only a single dimension, which is usually not the case for convolutional layers. Hence, the inputs and outputs of all layers in the equivalent network are the flattened versions of the original network.

Assume that all zeros tensor *W* has dimensions *M*×*N*×*C*×*M*×*N*×*L* where *M* and *N* are, respectively, the rows and columns of the input to a convolutional layer; *C* is the number of channels in the input; and *L* is the number of kernels in a layer. Additionally, assume *Kl* (the *l*-th kernel of the layer) has dimensions *H* × *W* × *C,* where H and W are the height and width of the kernel, respectively, and *C* is defined as before. We begin by populating *W* according to equation (3) for all values of *m, n,* and *l*. Then, we reshape *W* to (*M* * *N* * *C*) × (*M* * *N* * *L*) to obtain the 2D matrix that will transform the flattened input of the convolutional layer to the flattened output. Of course, *W* can be directly populated as a 2D matrix for improved performance.

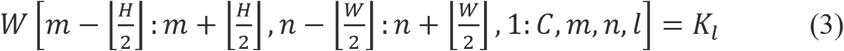

The calculation of LLRFs for the equivalent MLP network involves few steps (Fig 2B). The LLRF of a network with ReLU activations and no bias in the intermediate layers is equivalent to the gradient of the network’s output with respect to the input vector (81) (equations 4–5),

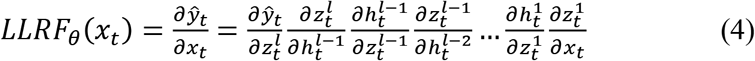

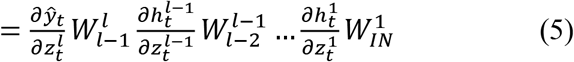

where 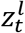 is the weighted sum of inputs to nodes in layer *l* for input *x_t_*, 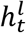, represents the output of nodes in layer *l* to the same input, and *θ* denotes the dependence on the parameters of the network. In a network with ReLU nodes:

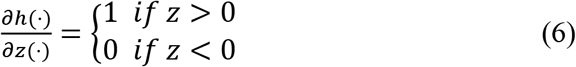

which means that we can replace the product of 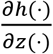 and 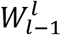 with an adjusted weight matrix 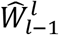 defined by equation (7), where *m* and *n* are indices of nodes in layer *l* and *l* — 1.

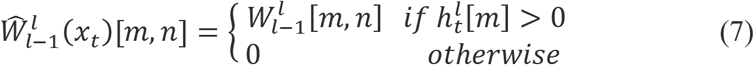

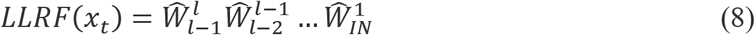

Because LLRF can be defined as the gradient of the output with respect to the input of the network, we can speed up the calculation process by utilizing TensorFlow’s built-in gradient calculation capability.

### Linearized function robustness

To ensure the validity of our LLRF analysis considering the intrinsic randomness involved in training deep neural networks using a stochastic gradient-descent algorithm from different parameter initializations by random sampling of training data, we need to show that our computed LLRFs are robust across different instances of a network. We do this by training 10 instances for each electrode, then grouping them into two equal sets of 5 networks each and averaging the LLRFs over instances for each group. We compare the LLRFs from the two groups for all time steps and electrodes (Supplementary Fig 3). The left panel histogram suggests that LLRF are robust to initialization, while the right panel shows that the similarity becomes even stronger when the LLRF is less noisy (has higher gain). To counter initialization noise, all LLRFs used in this paper are averaged over at least five network instances.

### Complexity estimation

To quantify the nonlinearity of the network receptive field, we measure the diversity of the equivalent linear functions that the network learns for different instances of the stimulus. To measure this function diversity, we calculated the singular-value decomposition (90) of the matrix containing all the linearized equivalent functions of a network (Fig 2D). Each singular value indicates the variance in its corresponding dimension; therefore, the steepness of the sorted singular values is inversely proportional to the diversity of the functions that are learned by the network. We define the complexity of the network as the sum of the normalized singular values(equation 9) where *σ_i_* is the *i*-th element of the singular values vector, and *D* is the length of the vector.

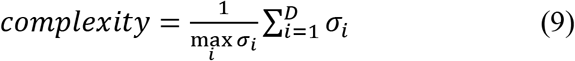

### Estimating gain change parameter

We calculated the magnitude of the LLRF at each time point using its standard deviation (equation 10, F = 32, T = 40). The gain change parameter for each site was then defined as the standard deviation of the LLRF magnitude over the duration of the test stimulus (equation 11). This quantity measures the degree to which the magnitude of the LLRF changes across different instances of the stimuli.

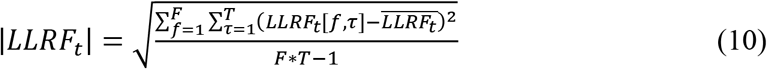

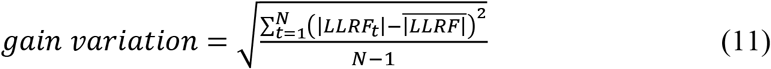

### Estimating temporal hold

Calculation of the temporal hold parameter involves multiple steps. For the LLRF at every time point, we calculated the number of future time steps in which the shifted LLRF functions were significantly correlated with the LLRF at the current time point (up to a maximum of 20 future time points corresponding to 200 ms, which was empirically determined as a sufficient upper limit). To accomplish this goal, we first zero padded the LLRFs by 200 ms on each side and shifted the LLRF at the *i_th_* future frame relative to the current frame by *i* time steps. The number of future frames for which we found a significant correlation between the current LLRF and future latency-shifted LLRFs indicates the number of time points in which the current LLRF feature will appear as shifted in latency, which we refer to as the temporal hold value of the LLRF. This procedure is shown in Supplementary Fig 5.

### Estimating shape change

The shape change parameter represents the diversity of the linear functions that is not due to the gain change or temporal hold nonlinearities. To calculate this parameter, we first normalized the effects of gain change and temporal hold by dividing the LLRFs by their magnitude and time aligning the shifted LLRF features. Finally, we repeated the same procedure used for the calculation of complexity equation (9) but instead used the normalized, shift-corrected LLRFs to calculate the singular-value decomposition. The sum of the sorted normalized singular values indicates the diversity of the linear functions learned by the network due to a change in their spectrotemporal feature tuning.

## Data availability

The data that support the findings of this study are available upon request from the corresponding author [NM].

## Code availability

The codes for pre-processing the ECoG signals and calculating the high-gamma envelope are available at *http://naplah.ee.columhia.edu/naplih.html* (91).

## Acknowledgments

This work was funded by a grant from the National Institutes of Health, NIDCD-DC014279, National Science Foundation CAREER award, and the Pew Charitable Trusts, Pew Biomedical Scholars Program.

## Author contributions

B.K. and N.M designed the experiment. B.K., M.K., J.H., N.M., A.D.M recorded the neural data. M.K., H.A., B.K., N.M. analyzed the data. M.K., N.M wrote the manuscript. All authors commented on the manuscript

