## Supplemental Figures and Video Captions for "Estimating and interpreting nonlinear receptive fields of sensory responses with deep neural network models"

**This PDF file includes:**

Supplementary Figures 1 to 9
Captions for Supplementary Videos 1 to 3

**Other Supplementary Materials for this manuscript include the following:**

Supplementary Videos 1 to 3

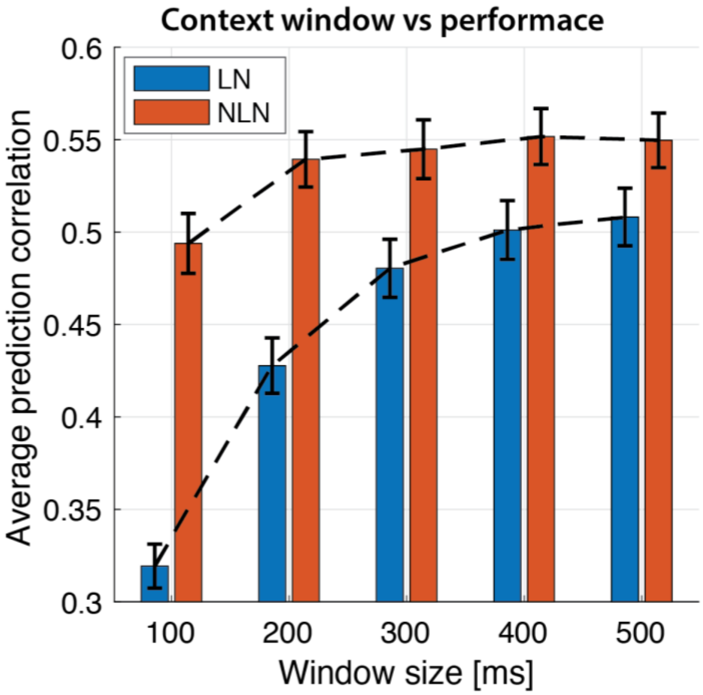


**Supplementary figure 1. Selecting stimulus window length for prediction**

Prediction accuracy of neural responses with varying the duration of the sliding window for LN and NLN models.

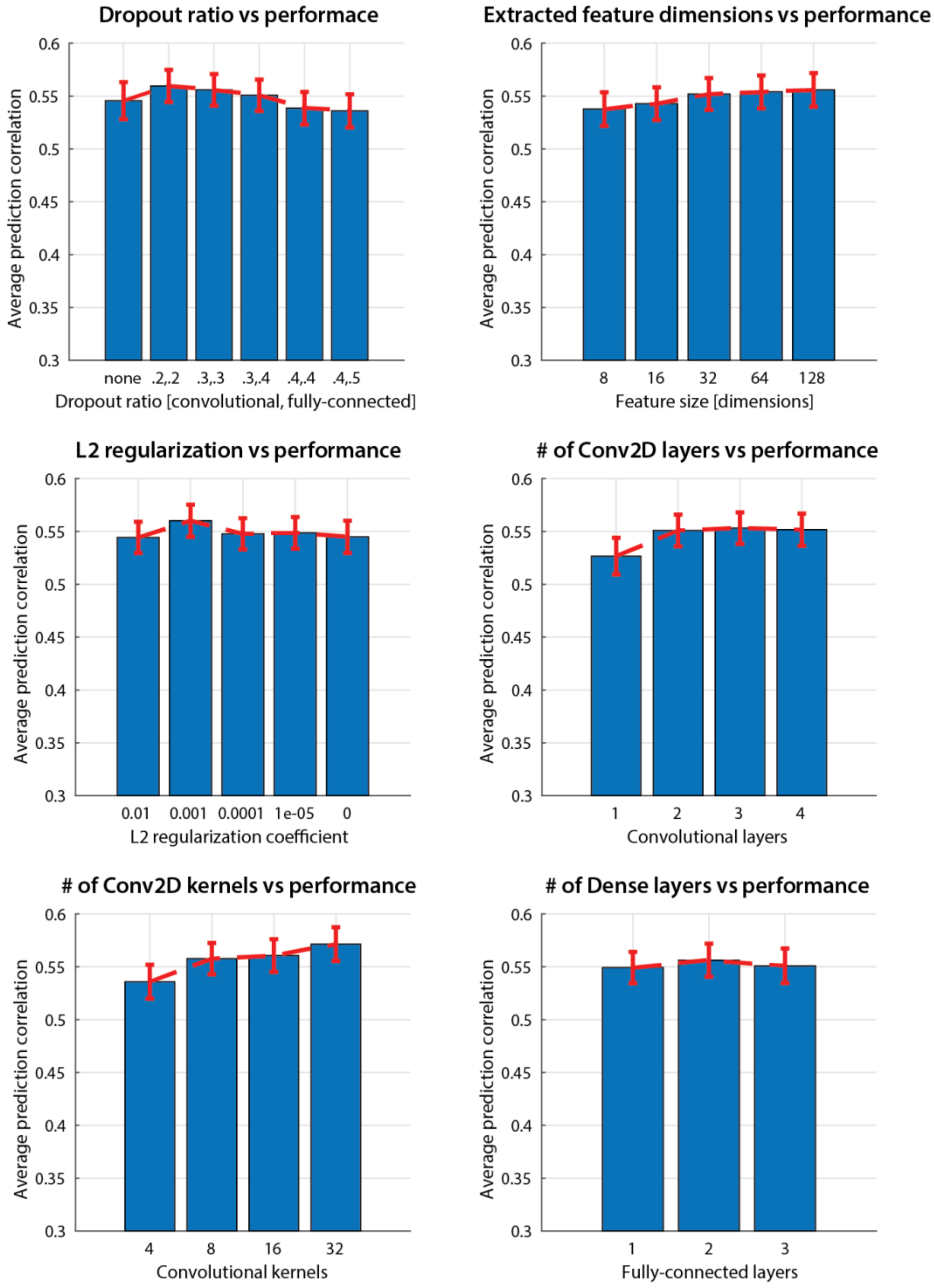


**Supplementary figure 2. Hyperparameter optimization**

Choosing the hyperparameters of the network to maximize prediction accuracy of the NLN model.

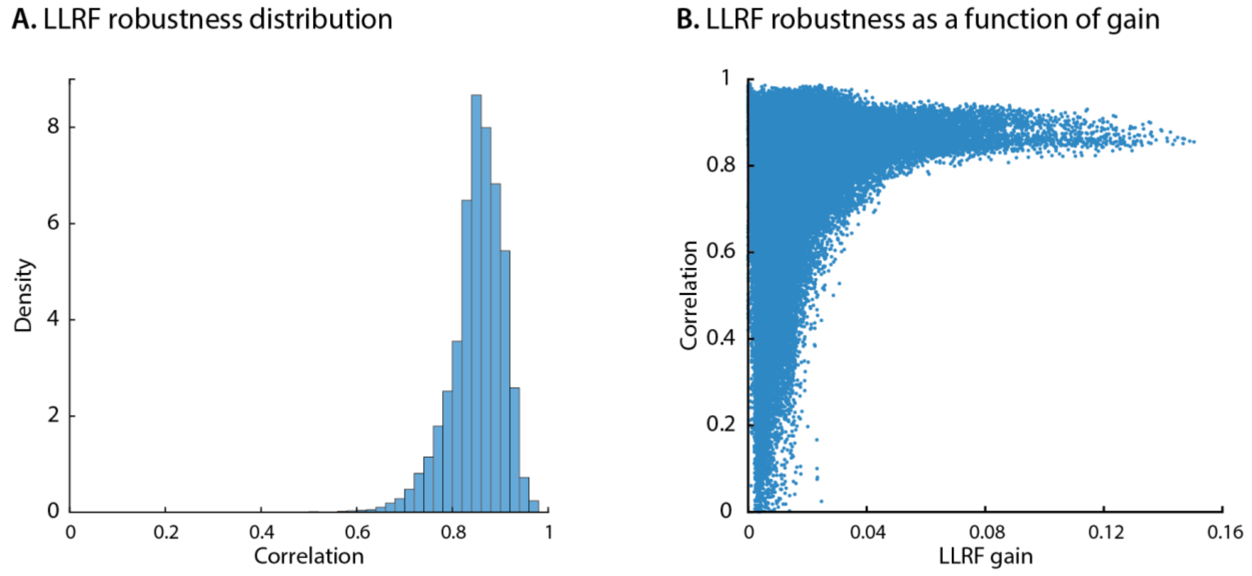


**Supplementary figure 3. LLRF robustness across initializations**

We trained 10 instances of the NLN model for each electrode and grouped them into two groups of evens and odds. For all time points we calculated the 2D correlation of the average LLRF from the even models with the average LLRF from the odd models. (A) a histogram of all values where one data point corresponds to the robustness for a specific electrode at a specific time point. (B) the relation of robustness with the gain of the linearized function. When the LLRF has higher gain the function is more robust.

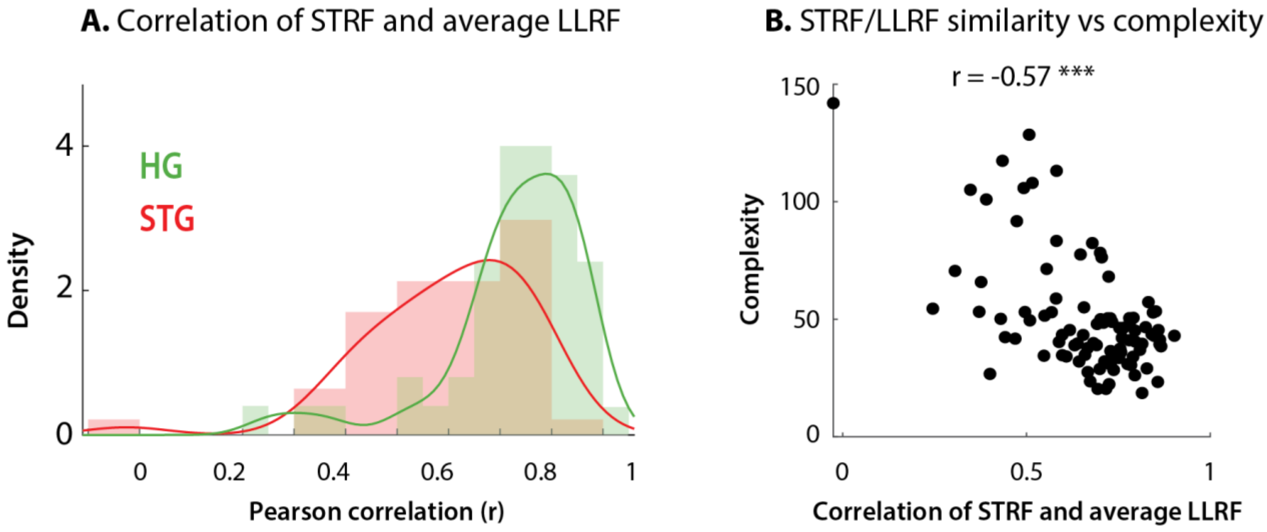


**Supplementary figure 4. Comparing LLRFs to STRFs**

(A) The average LLRF calculated across the test dataset is highly correlated with the calculated STRF, especially for HG electrodes. (B) Similarity of STRF and the average LLRF is inversely correlated with the complexity of the function the NLN model has learned, as expected.

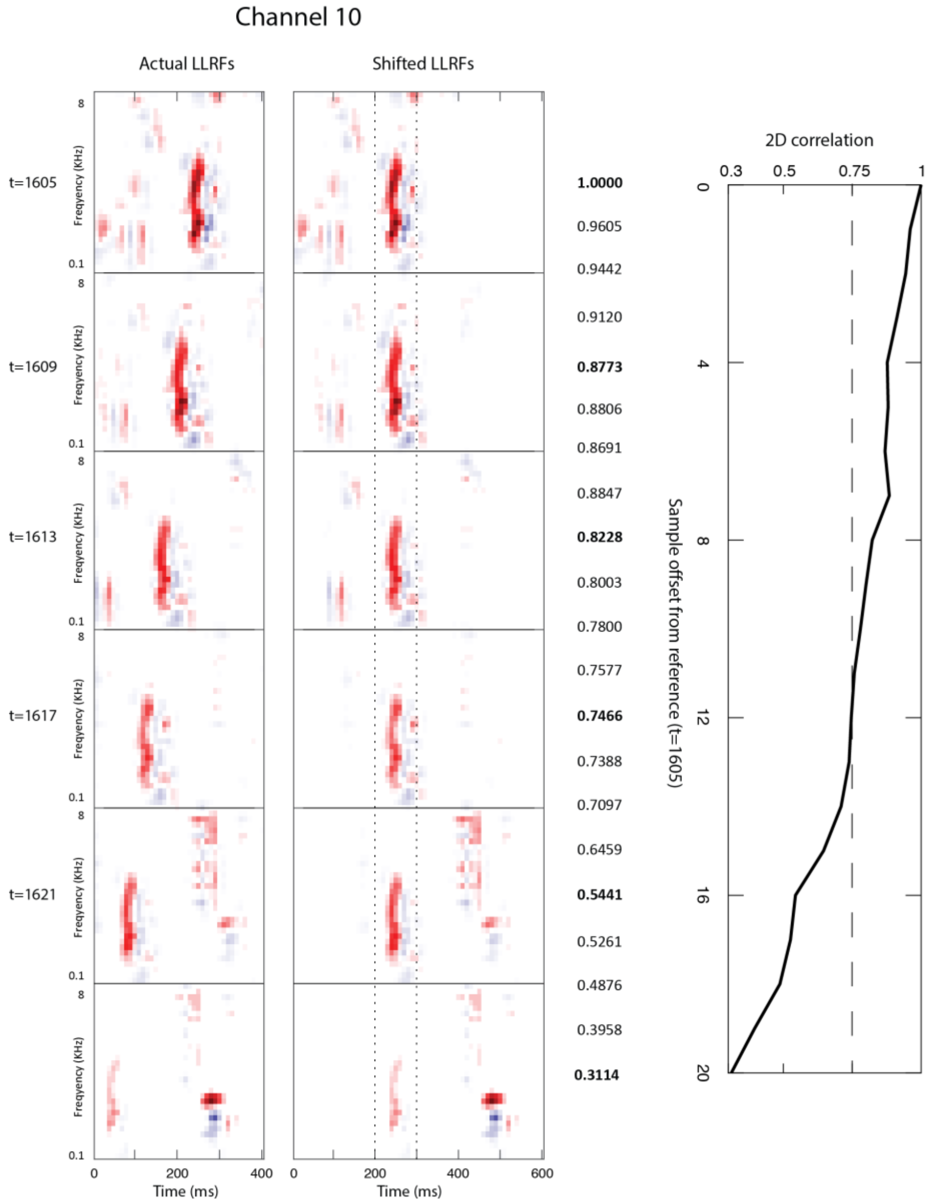


**Supplementary figure 5. Calculating temporal hold**

The leftmost LLRFs are the raw functions extracted from the network. The same spectrotemporal feature appears in all; however, the latency of this feature is shifted by one sample in every consecutive frame. The LLRFs on the right are the same weights but shifted by the duration of the distance of the time frames. The LLRFs in the second column are zero padded to allow shift operation. The cross correlation between the LLRF at each time point and the shifted LLRFs in future frames are used to assess the temporal hold property. The third column shows the correlation between the LLRF at t = 1605 and the next 20 shift-corrected LLRFs. The temporal hold value in this case is 110 ms because the next 11 shift-corrected LLRFs are significantly correlated with the LLRF at t = 1605.

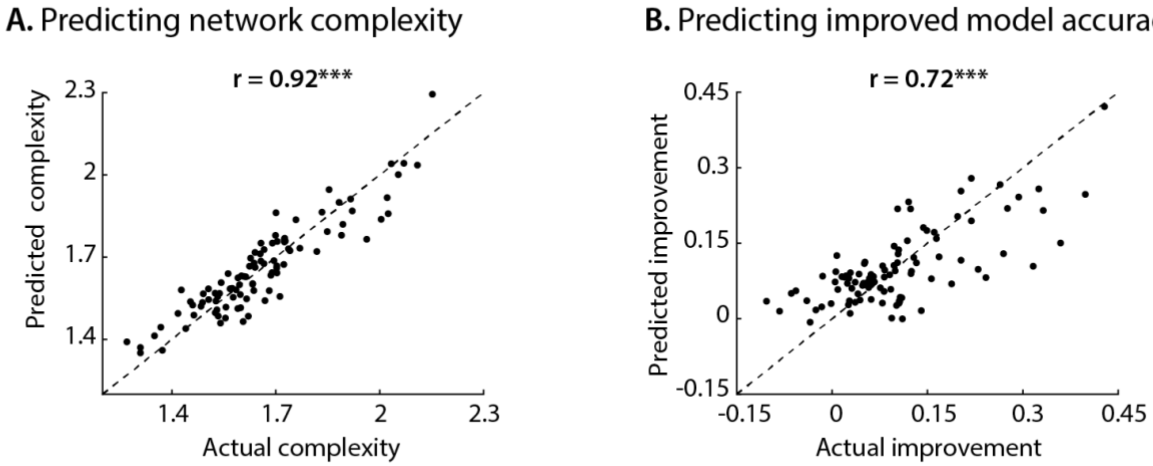


**Supplementary figure 6. Complexity and accuracy improvement predicted values**

True and predicted values for network complexity and improved model accuracy from the three nonlinear parameters.

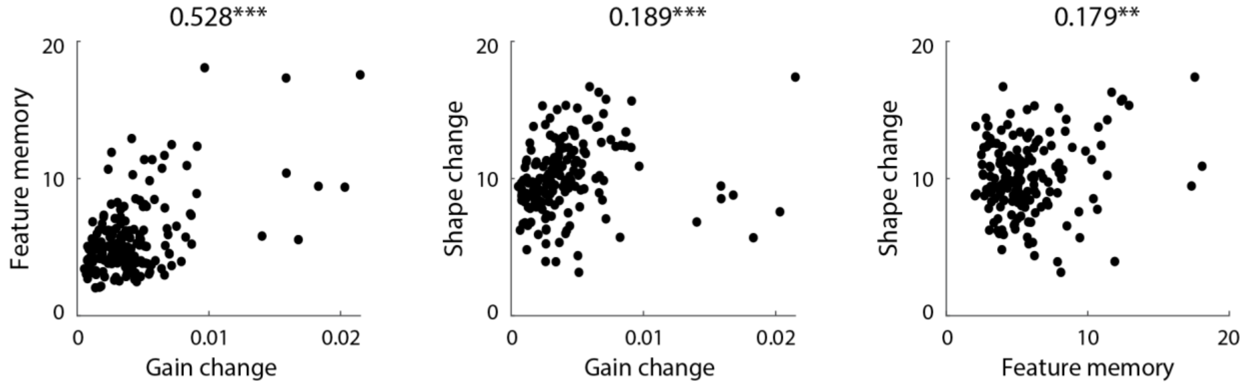


**Supplementary figure 7. Covarying nonlinear parameters**

The three parameters characterizing the nonlinear properties of neural sites (gain change, temporal hold, and shape change) are not independent and covary significantly.

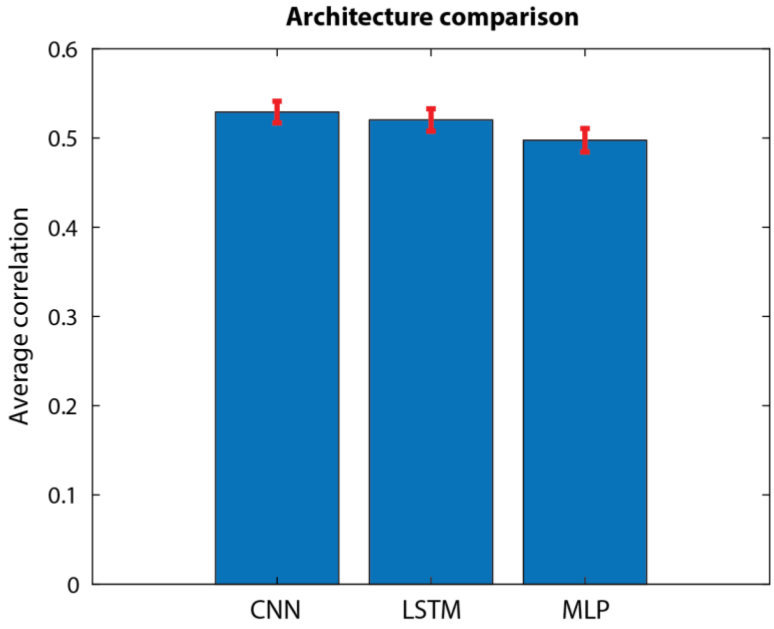


**Supplementary figure 8. Comparing with other DNN architectures**

Comparison of convolutional neural networks (CNN), long short-term memory networks (LSTM) and multilayer perceptron (MLP) in predicting the neural responses of all 99 electrodes used in the paper. The error bars represent the standard error.

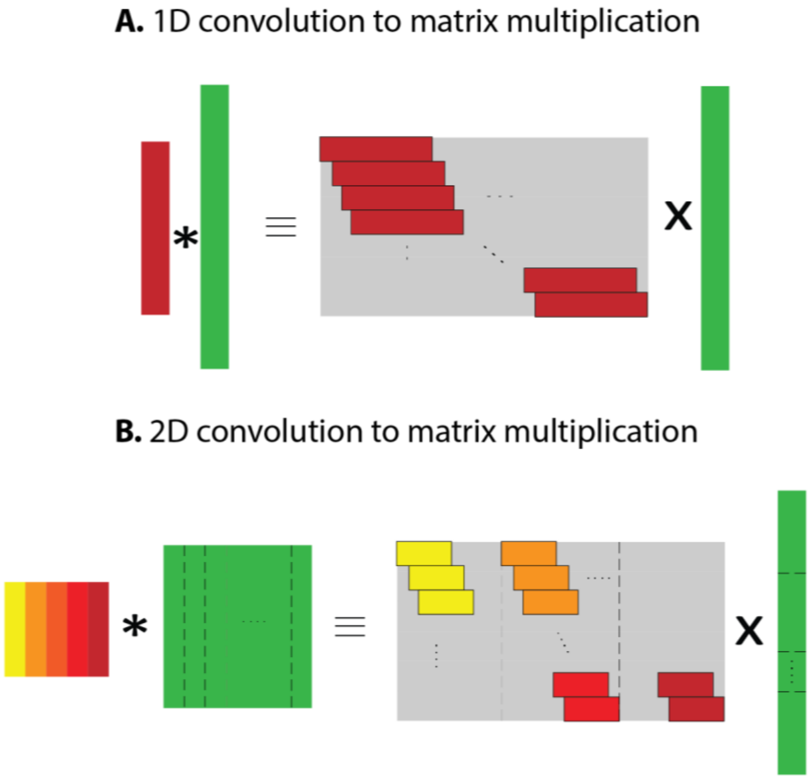


**Supplementary figure 9. Converting convolution to matrix multiplication**

Converting the convolutional neural networks into a feedforward network to simplify LLRF calculation. The convolution operation is linear; hence, it can be converted into a matrix multiplication. (a) Converting 1-d convolution into a 2-d matrix multiplication. The matrices are the shifted version of the 1-d vectors. The i-th row in the output will be the product of the input vectors (shown in green) and the convolution kernel (red) shifted by $i$ steps. (b) Converting 2-d convolution to a 2-d weight matrix using the same principle by flattening both the kernel and the input into one-dimensional forms by stacking the columns vertically on top of each other.

### Supplementary Videos

The following files can also be accessed online at http://naplab.ee.columbia.edu/sdlt.html

##### Supplementary Video 1

The spectrotemporal receptive field (STRF) model applies a weight function to the stimulus to predict the neural response.

##### Supplementary Video 2

The locally linear receptive field (LLRF) applies a time-varying, stimulus-dependent weight function to the stimulus to predict the neural response.

##### Supplementary Video 3

Visualization of gain change, temporal hold, and shape change for three example electrodes, each demonstrating one type of nonlinearity more prominently.
